# Membrane potential regulates Hedgehog signaling and compartment boundary maintenance in the *Drosophila* wing disc

**DOI:** 10.1101/2020.06.02.128892

**Authors:** Maya Emmons-Bell, Riku Yasutomi, Iswar K. Hariharan

## Abstract

The *Drosophila* wing imaginal disc is composed of two lineage-restricted populations of cells separated by a smooth boundary. Hedgehog (Hh) from posterior cells activates a signaling pathway in anterior cells near the boundary which is necessary for boundary maintenance. Here, we show that membrane potential is patterned in the wing disc. Anterior cells near the boundary, where Hh signaling is most active, are more depolarized than posterior cells across the boundary. Elevated expression of the ENaC channel Ripped Pocket (Rpk), observed in these anterior cells, requires Hh. Antagonizing Rpk reduces depolarization and disrupts the compartment boundary. Using genetic and optogenetic manipulations, we show that membrane depolarization promotes membrane localization of Smoothened and augments Hh signaling. Thus, membrane depolarization and Hh-dependent signaling mutually reinforce each other in this region. Finally, clones of depolarized cells survive preferentially in the anterior compartment and clones of hyperpolarized cells survive preferentially in the posterior compartment.

## Introduction

During the development of many organisms, boundaries between different groups of cells function as developmental organizers, serving as sources of morphogens, molecules that diffuse from their source and specify cell fates in a dose-dependent manner (reviewed by (Blair, 2003; Irvine & Rauskolb, 2001)). One of the best studied examples of this phenomenon is the anteroposterior (A-P) compartment boundary in the wing-imaginal disc of *Drosophila* (Garcia-Bellido et al., 1973). The wing disc arises from a population of approximately 20-30 embryonic cells in the second thoracic segment that straddle the parasegment boundary (Madhavan and Schneiderman, 1977; Worley, et al., 2013; Requena et al., 2017). These cells represent the primordium of the adult wing and the thoracic tissue to which it is attached.

During the larval stage of development, these cells proliferate to generate a structure, the wing imaginal disc, that is composed of approximately 40,000 cells (Martin et al., 2009). Throughout the course of organ growth and patterning, cells anterior to the parasegment boundary (A cells) and those posterior to the boundary (P cells) remain segregated as two lineage-restricted compartments (Blair, 2003). The boundary between the A and P compartments, the A-P compartment boundary, remains relatively smooth, indicating that these two populations of cells rarely intermingle. The mechanisms that separate these two populations of cells are not well understood (Dahmann et al., 2011; Batlle & Wilkinson, 2012). It has been suggested that differential adhesion might be the cause of this segregation, but compartment-specific adhesion molecules that could underlie this phenomenon have not been identified. There is increasing appreciation that mechanical forces function to keep these populations of cells separate. Indeed, during the later stages of larval development, the A-P boundary appears to be reinforced by an intracellular actomyosin cable that runs within cells that abut the boundary (Umetsu et al., 2014).

The segregation of the A and P cells enables the compartment boundary to act as an organizing center. P cells make the morphogen Hedgehog (Hh), which binds to its receptor Patched (Ptc), which is expressed exclusively in A cells. Hh has a short range either because of its limited diffusion or because it is taken up by nearby target cells via processes known as cytonemes. Thus, Hh alleviates the repressive effect of Ptc on the seven-transmembrane protein Smoothened (Smo) in A cells near the boundary, initiating a signaling cascade that culminates in the stabilization of the activator form of the transcription factor Cubitus interruptus (Ci), and expression of target genes such as the long-range morphogen Dpp (Jiang & Hui, 2008; Lee et al., 2016; Petrov et al., 2017). In turn, Dpp regulates imaginal disc patterning and growth in both compartments (Hamaratoglu et al., 2014).

While the role of cell-cell interactions, diffusible morphogens and even mechanical forces have been studied in regulating the growth and patterning of the wing disc, relatively little attention has been paid to another cellular parameter, membrane potential, or *V_mem_. V_mem_* is determined by the relative concentrations of different species of ions across the cell membrane, as well as the permeability of the membrane to each of these ions. These parameters are influenced by the abundance and permeability of ion channels, the activity of pumps, and gap junctions. While changes in *V_mem_* have been studied most extensively in excitable cells, there is increasing evidence that the *V_mem_* of all cells, including epithelial cells, can vary depending on cell-cycle status and differentiation status (Blackiston et al., 2009; Sundelacruz et al., 2009). Mutations in genes encoding ion channels in humans (“channelopathies”) can result in congenital malformations (Plaster et al., 2001). Similarly, experimental manipulation of ion channel permeability can cause developmental abnormalities in mice as well as in *Drosophila* (Dahal et al., 2017; Belus et al., 2018). Only more recently has evidence emerged that *V_mem_* can be patterned during normal development. Using fluorescent reporters of membrane potential, it has been shown that specific cells during *Xenopus* gastrulation and *Drosophila* oogenesis appear more depolarized than neighboring cells (Krüger & Bohrmann, 2015; Pai et al., 2015). A recent study established that cells in the vertebrate limb mesenchyme become more depolarized as they differentiate into chondrocytes, and that this depolarization is essential for the expression of genes necessary for chondrocyte fate (Atsuta et al., 2019). However, in many of these cases, the relationship between changes in *V_mem_* and specific pathways that regulate developmental patterning have not been established.

Here we investigate the patterning of *V_mem_* during wing disc development and show that the regulation of *V_mem_* has an important role in maintaining the compartment boundary and in regulating Hh signaling. We show that the cells immediately anterior the compartment boundary, a zone of active Hh signaling, are more depolarized than surrounding cells, and that Hh signaling and depolarized *V_mem_* mutually reinforce each other. This results in an abrupt change in *V_mem_* at the compartment boundary which appears necessary for normal boundary function. We also demonstrate that alterations in *V_mem_* regulate cell survival in a compartment-specific manner.

## Results

We began by asking whether or not *V_mem_* is patterned in the wing imaginal disc. Wing imaginal discs from third-instar larvae were dissected and incubated in Schneider’s medium containing the *V_mem_* reporting dye DiBAC_4_(3) (hereafter DiBAC) at a concentration of 1.9 μM for 10 min. DiBAC is an anionic, membrane-permeable, fluorescent molecule that accumulates preferentially in cells which are relatively depolarized compared to surrounding cells due to its negative charge, and has been used to investigate patterns of endogenous *V_mem_* in non-excitable cells in a variety of organisms (Adams & Levin, 2012; Krüger & Bohrmann, 2015; Atsuta et al., 2019). In contrast to patch clamp electrophysiology, utilizing DiBAC allowed us to make comparisons of membrane potential across a field of thousands of cells.

A dorsoventral stripe of cells running through the pouch of the wing disc appeared more fluorescent, thus indicating increased DiBAC uptake **(Fig. 1A, B)**, suggesting that these cells are more depolarized than surrounding cells. This pattern of fluorescence was observed in more than 35 individual wing imaginal discs, and was not observed when imaginal discs were cultured in the voltage-insensitive membrane dye FM4-64 **(Fig. 1C-E)**. Patterned DiBAC fluorescence was observed at both apical **(Fig. 1A-A’)** and basal **(Fig. 1B-B’)** focal planes. Addition of the Na^+^/K^+^ ATPase inhibitor ouabain to the cultured discs, which depolarizes cells by collapsing transmembrane Na^+^ and K^+^ gradients, resulted in increased, and more uniform, DiBAC fluorescence **(Fig. 1 F-G)**, indicating that patterned DiBAC fluorescence in wing disc tissue was contingent upon mechanisms that normally maintain *V_mem_*.

**Figure1.**
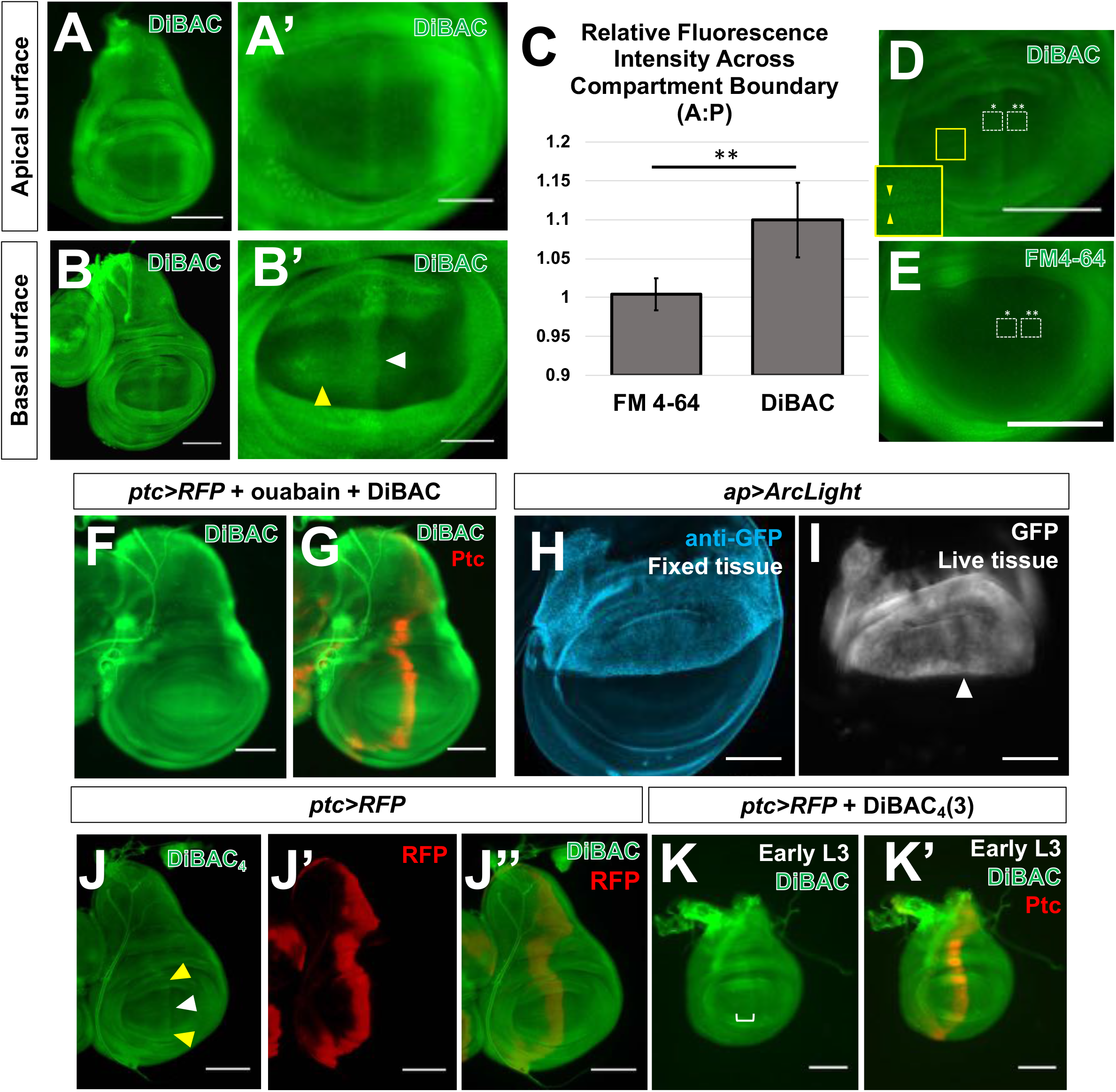
Membrane potential is patterned in the third instar wing-imaginal disc. (**A-B**) Live third instar discs incubated in DiBAC. DiBAC fluorescence is observed in the wing imaginal disc in both apical (**A-A’**), and basal (**B-B’**) optical sections. Increased fluorescence is also observed at the dorsoventral (D-V) compartment boundary in the anterior compartment (yellow arrow, **B’**, and inset, **D**). (**C-E**) Comparison of DiBAC with the voltage-insensitive dye FM4-64. Incubation of live discs in the voltage-insensitive pan-membrane dye FM4-64 (**E**) shows more uniform fluorescence when compared to DiBAC (**D**). Quantitative comparison of fluorescence in the two white boxes in each panel is shown in (**C**). N=7 discs for each treatment, ** indicates p<0.001 The region boxed in yellow in (**D**) is shown at higher magnification to show DiBAC fluorescence at the D-V compartment boundary. (**F, G**) Incubation in ouabain results in brighter and more uniform DiBAC fluorescence. **(H, I)** Discs expressing the genetically encoded membrane potential sensor ArcLight. Live discs (**I**) show a dorsoventral stripe of cells with decreased fluorescence, indicative of a more depolarized membrane potential (arrowhead). In fixed discs (**H**), relatively uniform expression of the indicator was observed using anti-GFP immunostaining. (**J, J’**) Live wing discs expressing *UAS-RFP* anterior to the compartment boundary, under the control of *ptc-Gal4*, were incubated in DiBAC, showing that the stripe of increased DiBAC fluorescence coincides with the posterior edge of *ptc* expression. White arrowhead in (**J**) indicates the stripe in the wing pouch; yellow arrowheads indicate the stripe in the dorsal and ventral hinge. (**K, K’**) Early L3 discs incubated in DiBAC. Patterned depolarization is evident throughout the third larval instar, with the stripe of increased fluorescence becoming narrower in more mature discs with developmental time (Compare **K** with **A**). Scale bars are 100μM in all panels, except for (**A’**, **B’**, **H**, and **I**), where scale bars are 50μM.

To determine if similar changes in patterned *V_mem_* could be visualized using an independent method, we used a genetically-encoded sensor of *V_mem_* ArcLight. ArcLight consists of a fusion of the S4 domain of a *Ciona intestinalis* voltage-sensitive phosphatase to a GFP variant (Jin et al., 2012; Cao et al., 2013). Upon depolarization, conformation of the voltage-sensitive domain is altered, leading to decreased GFP fluorescence. Therefore, depolarized cells appear less fluorescent than cells that maintain their resting potential. We expressed *UAS-ArcLight* in the dorsal compartment of the wing disc using *apterous-Gal4* (*ap-Gal4*). Using an anti-GFP antibody in fixed tissue, we observed uniform ArcLight expression in the *ap-Gal4* expression domain **(Fig. 1H)**. When GFP fluorescence was visualized in live discs, a dorsoventral stripe of cells appeared less fluorescent, indicating that they are more depolarized than surrounding tissue **(Fig. 1I)**. Thus, both DiBAC and ArcLight reveal similar patterns of depolarization in the third instar wing disc.

The position of the dorsoventral stripe of altered *V_mem_* is reminiscent of the anteroposterior (A-P) compartment boundary in the wing disc, which separates two lineage-restricted populations – the anterior (A) cells and the posterior (P) cells. In order to identify the population of depolarized cells with respect to the compartment boundary, we examined discs expressing *UAS-RFP* under the control of *patched-Gal4* (*ptc*>RFP) that had been incubated in DiBAC. *ptc*>RFP is expressed in those cells that express the highest levels of endogenous *ptc*, which are immediately anterior to the compartment boundary. The cells in which DiBAC accumulated at higher levels correlated with expression of RFP throughout the third larval instar **(Fig. 1J-J’’)**, indicating that cells anterior to the compartment boundary are more depolarized than cells across the compartment boundary in the posterior compartment. *ptc*-expressing cells in the hinge also were more fluorescent upon DiBAC staining **(Fig. 1J)**, but for the purposes of this work we focused on the wing pouch. As with *ptc-Gal4* expression, the domain of relative depolarization was broader in early third-instar wing discs, becoming more and more restricted to cells just anterior to the compartment boundary over the course of developmental time **(Fig. 1K-K’)**.

We examined other imaginal discs using DiBAC **(Figure 1, Figure Supplement 1)**. We did not detect increased fluorescence either at the compartment boundary of the leg disc or of the antennal disc. In the eye disc, we observed increased fluorescence in the region of the “ocellar spot”, the primordium of the ocelli, which also expresses *ptc>RFP*. Thus, it is possible that the altered depolarization at the compartment boundary might be specific to the wing disc. Alternatively, the changes in *V_mem_* in these other discs might be more subtle and not detectable using DiBAC. We therefore focused our efforts on studying the wing disc.

### Altered expression of endogenous ion channels anterior to the compartment boundary

The resting potential, *V_mem_*, results from the activity of a large number of different transporters of charged molecules, as well as the permeability of the membrane to each of those molecules. Thus, the relative depolarization of the region immediately anterior to the compartment boundary is unlikely to result simply from a change in the activity of a single pump or channel. However, by identifying transporters expressed in this region, it should be possible to manipulate *V_mem_* by altering their expression or properties. To that end, we examined a published transcriptome dataset (Willsey et al., 2016), comparing the abundance of transcripts in *ptc*-expressing cells with those of cells in the posterior compartment. In this dataset, we noticed several ion channels with differential expression between the two populations of cells. Among these are two members of the Degenerin Epithelial Na^+^ Channel (DEG/ENaC) family of channels, *ripped pocket* (*rpk*) (C. M. Adams et al., 1998), and *pickpocket 29* (*ppk 29*) (Thistle et al., 2012). DEG/ENaC channels are members of a diverse family of amiloride-sensitive cation channels. An antibody that recognizes the Rpk protein has been characterized previously (Hunter et al., 2014), which allowed us to examine its pattern of expression. In late L3 wing discs, we found that Rpk was indeed expressed anterior to the compartment boundary, in a stripe of cells that also express *ptc>RFP* **(Fig. 2A-A’)**. In addition, we observed expression of Rpk in cells near the dorsoventral (D-V) boundary. Expression was most obvious in two rows of cells flanking the D-V boundary in the anterior compartment, which are likely to be the two rows of cells arrested in the G2 phase of the cell-cycle (Johnston & Edgar, 1998). Indeed, the pattern of DiBAC uptake in this portion of the wing disc also suggests a very thin stripe of low-fluorescence flanked by two regions of higher fluorescence **(see inset in Fig. 1D)**. This is consistent with previous work showing that cells in culture become more depolarized as they progress through S-phase, and peaks at the onset of mitosis (Cone, 1969), reviewed in (Blackiston et al., 2009). Thus, at least in principle, the increased expression of Rpk could contribute to the depolarization observed in these regions of the wing disc.

**Figure 2.**
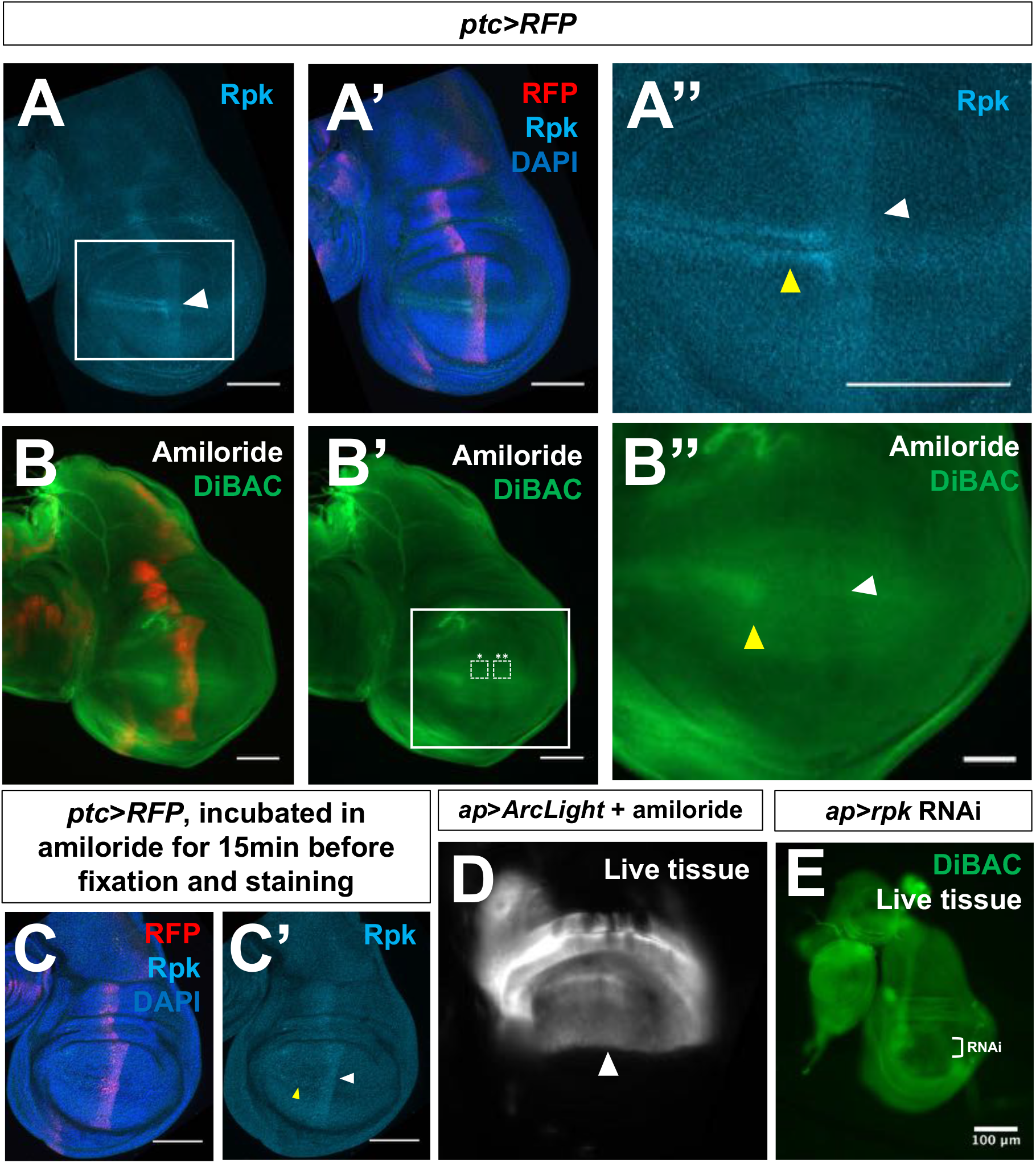
Expression of endogenous channels is patterned and contributes to depolarization anterior to the compartment boundary. (**A-A”**) Expression of Rpk is increased anterior to the A-P compartment boundary, and in two rows of cells flanking the D-V compartment boundary in the A compartment. (**B-B”**) Blockade of DEG/ENaC channels by incubation in amiloride abolishes the dorsoventral stripe of increased DiBAC fluorescence (white arrowhead). White boxes indicate ROIs used to calculate average DiBAC fluorescence intensity ratio across the compartment boundary = 0.99AU, s.d. 0.04, n = 4 discs. Increased fluorescence at the D-V boundary in the A compartment is still observable (yellow arrowhead). (**C, C’**) Amiloride incubation does not diminish Rpk expression. (**D**) Discs expressing ArcLight using *ap-Gal4* no longer have the dorsoventral stripe of reduced fluorescence shown in Figure 1G after amiloride treatment. (**E**) Expression of *rpk* RNAi in the dorsal compartment of the imaginal disc results in diminished DiBAC fluorescence. Patterns of DiBAC fluorescence in ventral tissue are challenging to interpret due to the altered morphology of discs of this genotype. Scale bars are 100μM, except in (**B’’**), where scale bars are 50μM.

**Figure 1, Figure Supplement 1.**
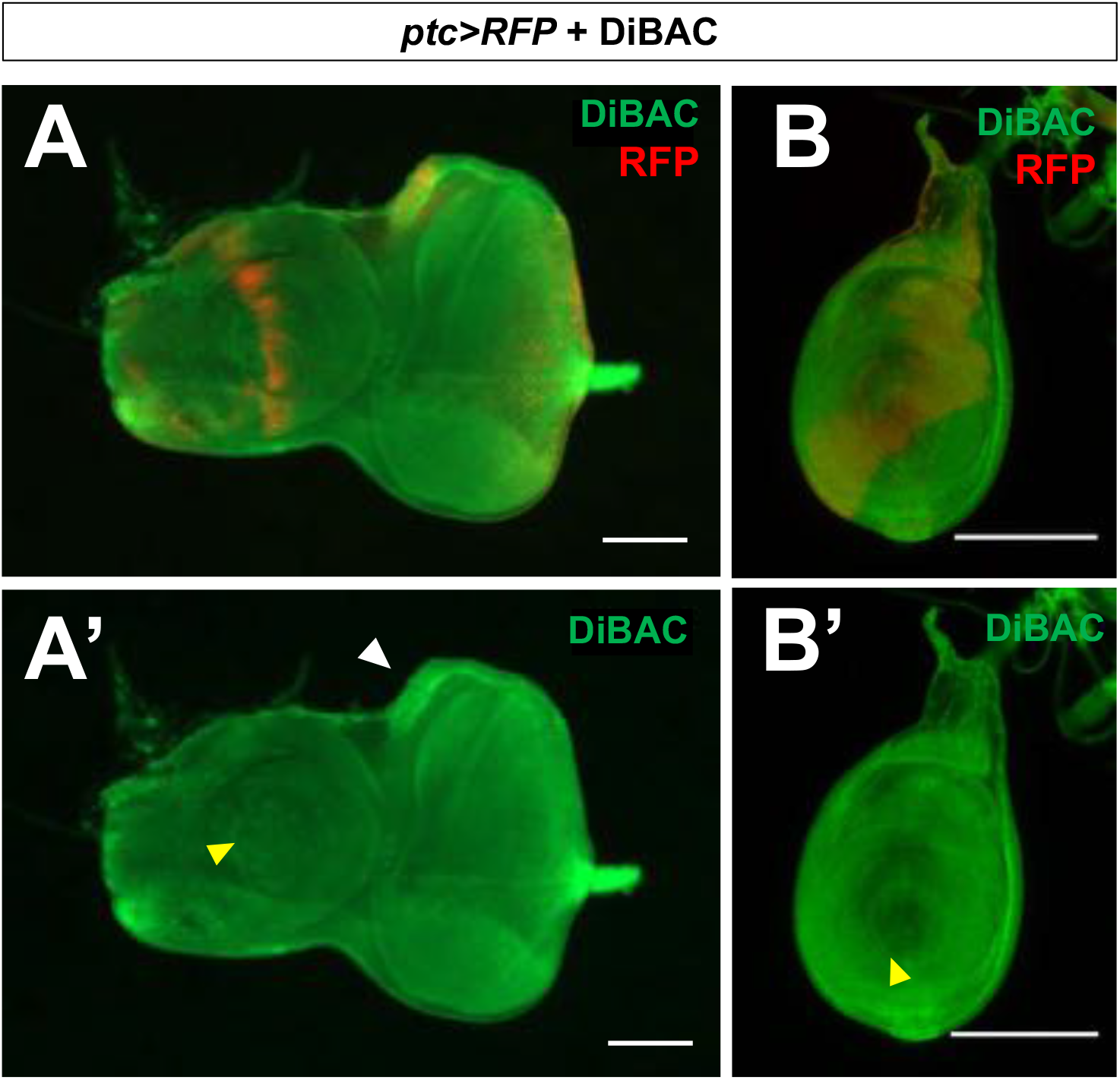
DiBAC fluorescence in eye-antennal and leg discs. (**A, A’**) Eye-antennal discs Increased fluorescence is observed in the region of the primordium of the ocelli which also expresses *ptc>RFP*. (**B, B’**) Leg imaginal discs. No change in DiBAC fluorescence is observed at the compartment boundary of the leg disc. Yellow arrowheads indicate compartment boundaries. Scale bars are 100μM in (A-A’) and 50μM in (B-B’). In each panel, discs are positioned such that the anterior is to the left, and dorsal is up.

Since Rpk, and possibly Ppk29, are expressed anterior to the compartment boundary, we tested the effect blocking these channels by treating discs with amiloride. Amiloride is predicted to reduce the permeability of DEG/ENaC-family channels (Garty, 1994). Addition of amiloride did not alter the pattern of Rpk protein expression **(Fig. 2C-C’)**. However, amiloride addition abolished the stripe of increased DiBAC fluorescence anterior to the compartment boundary **(Fig. 2B-B’’)**, as well as the stripe of decreased fluorescence observed with ArcLight **(Fig. 2D)**. These findings suggest that a conductance mediated by one or more channels of the DEG/ENaC family contributes to the relative depolarization of this region. Amiloride addition did not, however, seem to affect the increased DiBAC fluorescence observed at the D-V boundary **(Fig. 2B’’)**. Since amiloride likely targets multiple ENaC channels, we also depleted Rpk using an RNAi transgene expressed in the dorsal compartment of the disc using *ap-Gal4*. In these discs, we observed reduced DiBAC fluorescence in the dorsal portion of the disc **(Fig. 2E)**. However, due to altered disc morphology, it was difficult to assess whether the stripe of increased DiBAC fluorescence in the ventral portion of the disc was still observed. From these experiments, we conclude that reducing the expression or permeability of endogenous DEG/ENaC family channels, notably Rpk, can abolish the region of depolarization anterior to the A-P compartment boundary. Hence, Rpk, and possibly other DEG/ENaC channels, contribute to this local alteration in *V_mem_*.

In most cells, the Na^+^/K^+^ ATPase is primarily responsible for setting a negative *V_mem_*, since it uses ATP hydrolysis to extrude three Na^+^ ions and bring in two K^+^ ions per cycle of activity (Morth et al., 2007). RNA of *ATPα* (Lebovitz et al., 1989), which encodes a subunit of the Na^+^/K^+^ ATPase, was also detected at higher levels in *ptc*-expressing cells (Willsey et al., 2016). Using an antibody to ATPα, (Roy et al., 2013) we once again observed elevated expression anterior to the compartment boundary, with a hint of increased expression at the D-V boundary **(Fig. 3 A’’’)**. Additionally, RNA of Irk1, which encodes a K^+^ channel (MacLean et al., 2002), was reported to be decreased in *ptc*-expressing cells (Willsey et al., 2016). However, an antibody to this protein is currently not available. While it is difficult to predict the contribution of patterned expression of each channel to the patterning of *V_mem_*, detection of these channels in that portion of the wing disc allows us to manipulate their expression or to pharmacologically alter their properties.

**Figure 3.**
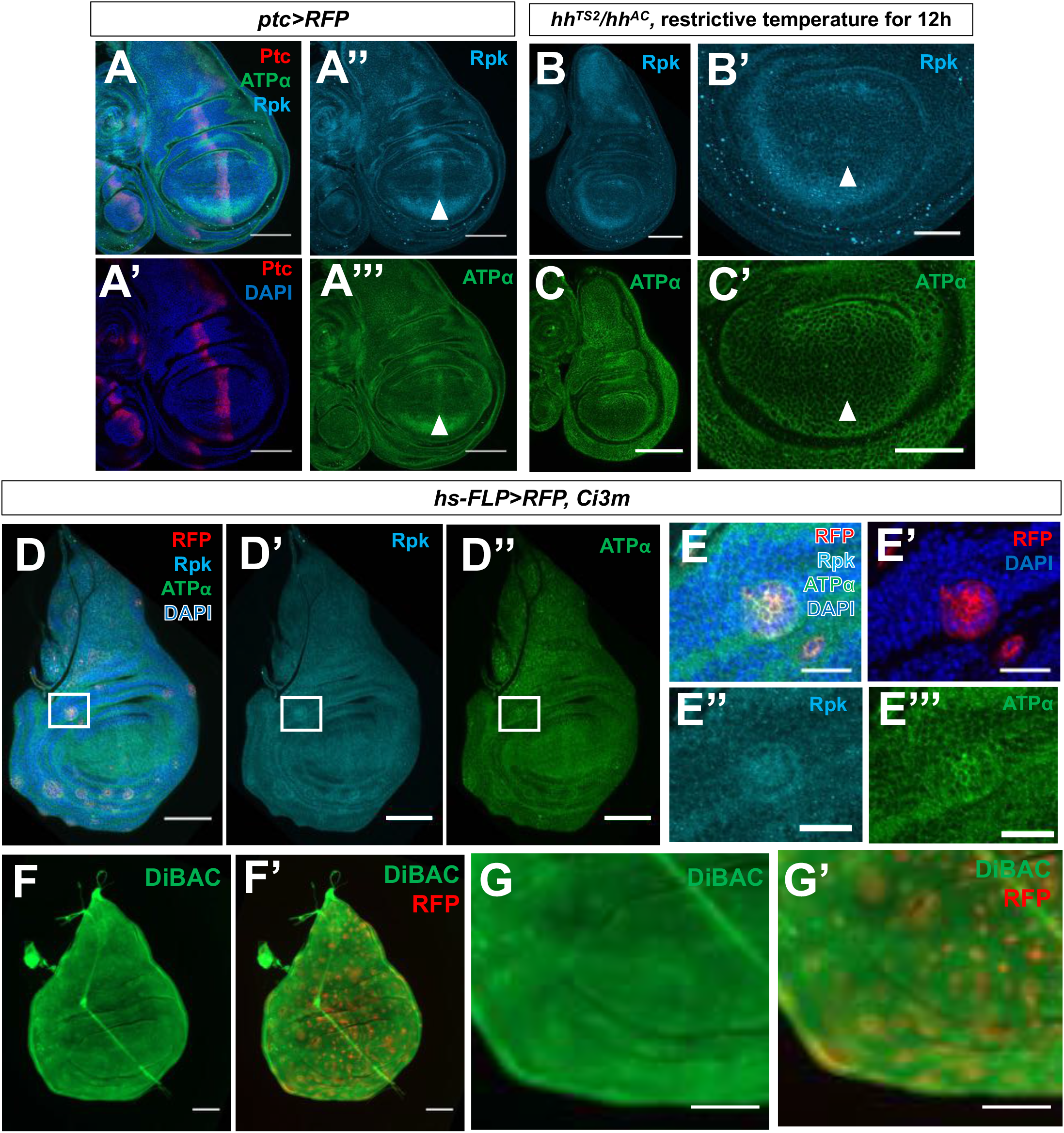
Patterned Rpk and ATPα expression and membrane depolarization require Hh signaling. (**A-A’”**) Immunostaining of discs expressing *ptc>RFP* with antibodies to rpk and ATPα showing elevated levels of both proteins anterior to the compartment boundary. (**B-C’**) L3 discs that are temperature-sensitive for *hh* following shift to the restrictive temperature for 12 h show loss of increased expression of Rpk (**B, B’**) and ATPα (**C, C’**) anterior to the compartment boundary. (**D-E’’’**) Clones of cells expressing a constitutively activated form of the transcription factor Ci show elevated rpk and ATPα expression. (**E-E’’’**) A single clone in the A compartment is shown at higher magnification. (**F-G’**) Live discs with clones of cells expressing activated Ci following incubation in DiBAC. The clones also express RFP. All scale bars are 100uM, except for (**B’’**), (**C’’**), and (**F-G’**) where scale bars are 50μM, and (**E-E’’’**), where scale bars are 25μM.

### Patterned expression of *rpk* and *ATPa* is regulated by Hedgehog signaling

Since the increased expression of Rpk and ATPα anterior to the compartment boundary occurs precisely within the region of increased Hh signaling **(Fig. 3A-A’’’)**, we tested whether manipulating components of Hh signaling pathway could alter expression of Rpk or ATPα. We used a temperature-sensitive *hh* allele (*hh^TS2^*) (Ma et al., 1993) in order to decrease Hh signaling for a short period of time. Larvae were raised at a permissive temperature (18°C) to permit normal *hh* function during early development, and then shifted to a restrictive temperature (30°C) during the third larval instar in order to reduce *hh* function, and dissected 12 hours after the temperature shift. Under these conditions, increased expression of either ATPα or Rpk was not observed anterior to the compartment boundary **(Fig. 3B-C’)**, indicating that normal levels of *hh* activity are necessary for the increased expression of these proteins anterior to the compartment boundary. Interestingly, some expression of Rpk is still visible near the D-V boundary in the anterior compartment which is likely *hh*-independent. We then examined the effects of increasing Hh signaling. Since Hh-dependent gene expression in this region typically results from stabilization of the activator form of Cubitus interruptus (Ci) (Aza-Blanc et al., 1997), we generated clones of cells expressing a constitutively active version of Ci (Ci3m (Price & Kalderon, 1999)). The Ci3m allele has S to A mutations at PKA-phosphorylation sites 1-3, rendering the protein more resistant to proteolytic cleavage. These clones had modest increases in expression of both of Rpk and ATPα **(Fig. 3D-E’’’)**. Additionally, these clones showed increased DiBAC fluorescence **(Fig. 3F-G’)**. Thus, Hh signaling appears to promote expression of both Rpk and ATPα, and relative depolarization.

### Manipulating expression of endogenous ion channels modulates Hedgehog signaling

We then investigated whether a depolarized *V_mem_* is required for the high levels of Hh signal transduction that occur immediately anterior to the compartment boundary. To that end, we reduced *rpk* expression in the dorsal compartment of the wing imaginal disc using *ap-Gal4* and *UAS-rpk^RNAi^*. The Hh activator Smo is transcribed throughout the *Drosophila* wing imaginal disc, but immunostaining for Smo protein reveals that it is most abundant on cell membranes in the posterior compartment, and in cells directly anterior to the compartment boundary **(Fig. 4A-A’’’)**. Membrane localization of Smo is thought to depend on high levels of Hh signaling (Zhu et al., 2003). The Hh signaling pathway is active in several rows of cells immediately anterior to the compartment boundary that receive Hh. Additionally, several components of the Hh signaling pathway are active in the entire posterior compartment, presumably because the absence of Ptc renders Smo constitutively active (Ramírez-Weber et al., 2000). However, target genes are not activated in posterior cells because Ci is only expressed in the anterior compartment. Knockdown of *rpk* resulted in a reduction in membrane staining of Smo in the dorsal compartment, as compared to ventral cells **(Fig. 4B’)**. This was observed in both posterior and anterior cells. In anterior cells near the compartment boundary, Hh signaling also results in stabilization of the activator form of Ci. In *ap>rpk^RNAi^* discs, the level of activated Ci in the dorsal compartment was reduced **(Fig. 4B’’)**. We also examined the expression of *ptc*, which is a direct transcriptional target of Ci (Alexandre et al., 1996). The stripe of staining with anti-Ptc was much fainter in the dorsal part of the disc **(Fig. 4C)**. Thus expression of *rpk^RNAi^* reduces Hh signaling in cells anterior to the compartment boundary.

**Figure 4.**
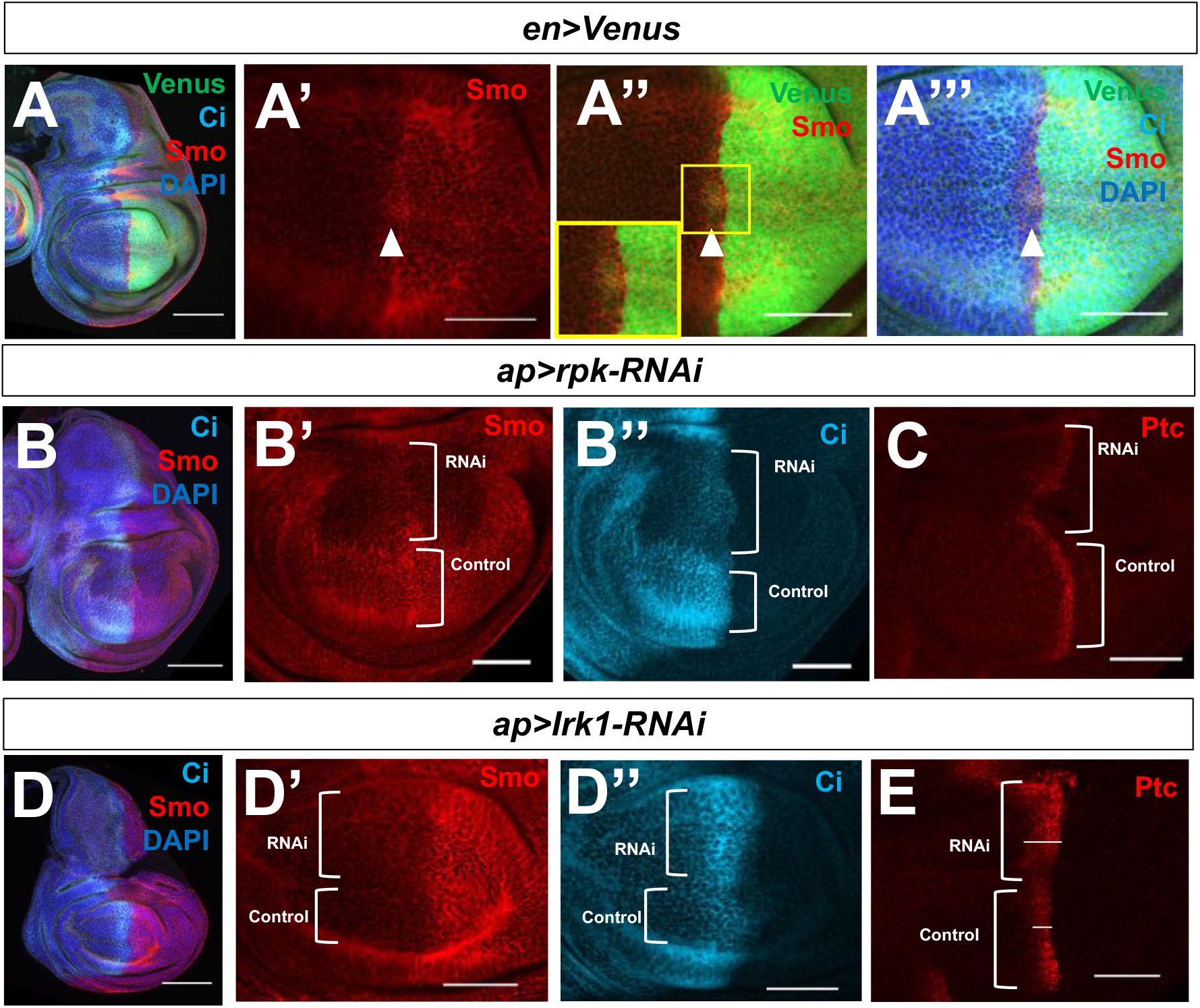
Manipulating levels of the ion channels Rpk and Irk1 impacts Hh signal transduction. (**A-A’”**) Immunostaining of Smo protein (red) and full-length Ci (light blue) in discs expressing *en-Venus*. Smo is expressed at higher levels in the P compartment and in a stripe 5-10 cells wide just anterior to the compartment boundary. (**B-E**) Effect of expressing an RNAi against the ENaC channel *rpk* (**B-C**) or the K^+^ channel *Irk1* (**D-E**) in the dorsal compartment of the disc using *ap-Gal4*. Knockdown of rpk in the dorsal compartment (**B-C**) results in decreased accumulation of the Hh activator Smo (**B, B’**), and the activator form of Ci (**B, B’’**). The expression of *ptc*, visualized using anti-Ptc (**C**), a downstream target gene of Hh signaling, is also diminished. Knockdown of Irk1 in the dorsal compartment results in increased accumulation Smo (**D, D’**), the active form of Ci (**D’’**), and Ptc (**E**). All scale bars are 50 μM, except for (**A)**, (**B**), and (**D**), where scale bars are 100μM.

Knockdown of *Irk1*, which encodes a K^+^ channel (MacLean et al., 2002), would be predicted to depolarize cells. Knockdown of *Irk1* in the dorsal compartment resulted in increased Smo membrane accumulation in the anterior compartment, an expansion of the domain of active Ci, and expanded Ptc expression **(Fig. 4D-E)**. Thus, manipulations that are predicted to increase depolarization increase Hh signaling, while those predicted to reduce depolarization reduce Hh signaling.

### Manipulating *V_mem_* regulates Smoothened localization

Cells of the wing imaginal disc are quite small and columnar, posing a challenge for subcellular imaging. Cells of the salivary gland are many hundreds of times larger than those of the wing disc, allowing for easier visualization of subcellular protein localization. Additionally, much work characterizing the regulation of Smo has been carried out in this tissue (Zhu et al., 2003). To directly test whether or not altering *V_mem_* can modulate Hh signaling, we used the bacterial sodium channel NaChBac, which can be used to cause membrane depolarization in insect cells by overexpression (Ren et al., 2001; Luan et al., 2006; Nitabach et al., 2006). Expression of NaChBac in the salivary gland using the Gal4 driver line *71B-Gal4* showed a clear increase in membrane-associated staining with anti-Smo **(Fig. 5A, B, D)** as well as increased expression of Ptc **(Fig.5 E-F)**. Correspondingly, increased expression of the mammalian potassium channel Kir2.1, which would be predicted to hyperpolarize *Drosophila* cells (Baines et al., 2001; Hodge, 2009), reduces fluorescence at the cell surface and increases intracellular fluorescence **(Fig. 5C, D)**. Thus sustained alteration in *V_mem_* modulates Hh signaling as assessed by Smo localization and Ptc expression.

**Figure 5.**
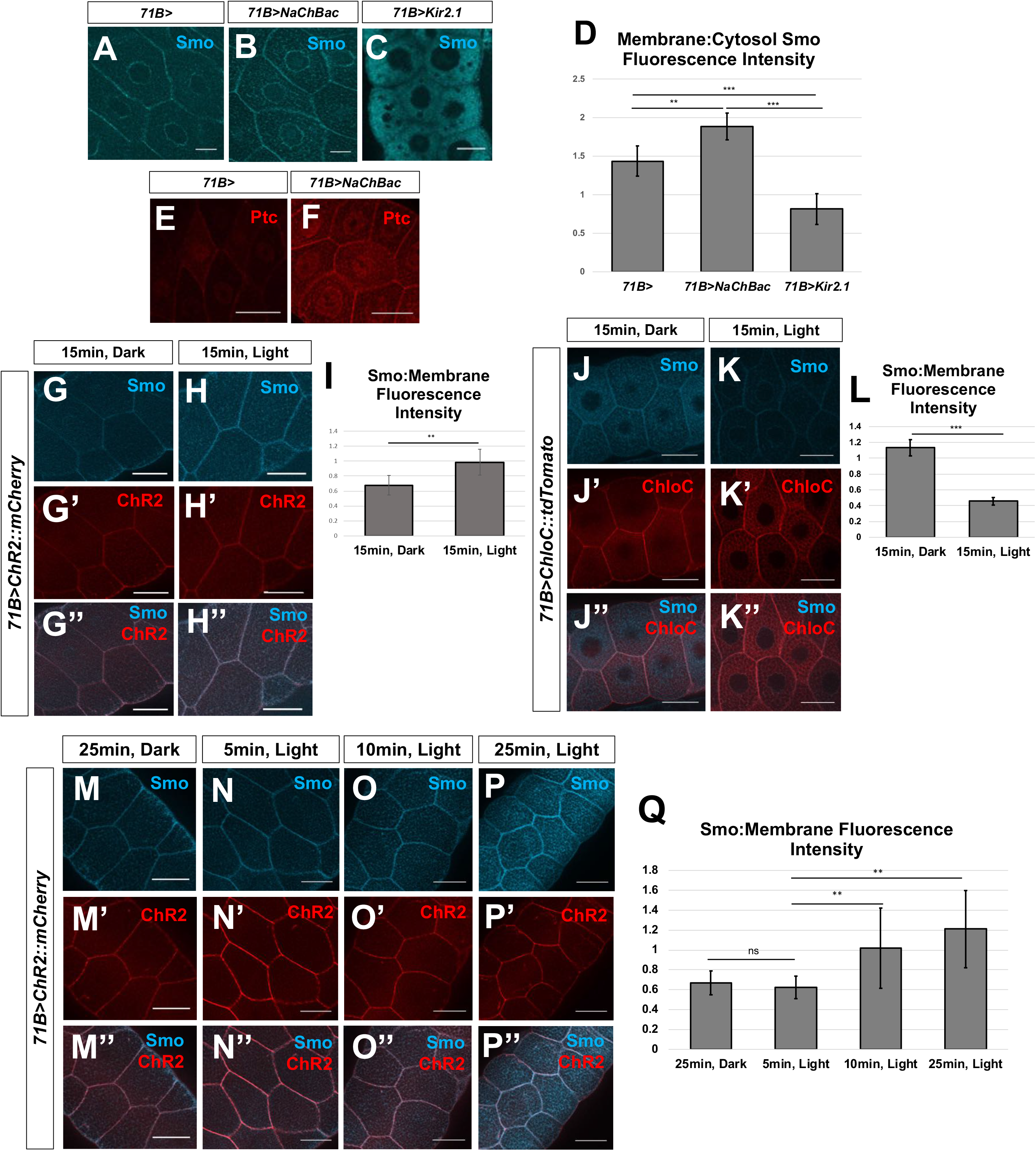
Membrane potential modulates Smo membrane abundance and downstream Hh signal transduction in larval salivary glands. (**A-Q**) Manipulation of *V_mem_* in salivary glands dissected from third-instar larvae. **(A-C)** Localization of Smo protein (blue) following expression of *71B-Gal4* (**A**), *71B-Gal4* and *UAS-NaChBac* (**B**) or *71B-Gal4* and *UAS-Kir2.1* (**C**). Expression of the sodium channel NaChBac increases membrane localization of Smo while expression of the potassium channel Kir2.1 reduces membrane localization, and increases cytosolic Smo (**B**, **C**). (**D**) Quantification of the ratio of membrane Smo fluorescence to cytosol fluorescence. N=7 salivary glands per genotype. Data were compared using ANOVA followed by Tukey test for significance (** indicates p<0.01, *** indicates p<0.001). (**E, F**) Expression of the Hh target gene *ptc* is visualized using anti-Ptc immunostaining. Compared to *71B-Gal4* (**E**), Ptc expression is increased in glands expressing *71B-Gal4* and *UAS-NaChBac* (**F**). (**G-I**) Effect of expression and activation of the depolarizing channelrhodopsin ChR2 on Smo protein membrane abundance. After dissection, glands were kept in either dark conditions (**G**) or exposed to 15 minutes of blue light (**H**). (**I**) Quantification of membrane fluorescence; N=13 glands, data compared using an unpaired t test. (** indicates p<0.01). (**J-L**) Effect of expression and activation of the Cl^-^-selective channelrhodopsin ChloC. An overall reduction in Smo immunostaining was observed following channel activation (**K**), as compared to control glands kept in the dark (**J**). (**L**) Quantification of membrane fluorescence; N=10 glands, data were compared using an unpaired t test, (*** indicates p<0.001). (**M-Q**) Time course of change in Smo localization. Salivary glands expressing the channelrhodopsin ChR2 were subjected to increasing intervals of activating light, then fixed and stained for Smo protein (**M-P”**). (**Q**) Quantification of membrane-associated fluorescence; N=7 glands/timepoint, data were compared using ANOVA followed by Tukey test for significance (** indicates p<0.01). Scale bars are 50μm, except for (**A-C**), where scale bars are 25μM.

In order to investigate the short-term consequences of altering *V_mem_* we used channelrhodopsin ChR2, which when exposed to blue light causes membrane depolarization (Nagel et al., 2003; Schroll et al., 2006). In addition to providing a second, independent way of depolarizing cells, this approach allowed us to examine short-term changes in Hh signaling that occur in response to depolarization. Dissected, ChR2-expressing salivary glands were either kept in darkness or exposed to activating light for variable intervals of time. Compared to glands kept in the dark, light-exposed glands had higher levels of Smo at the cell membrane **(Fig. 5G-I)**, suggesting that even relatively short-term depolarization can facilitate Smo accumulation at the cell surface. A time course showed that an increase in Smo membrane abundance was detectable as early as 10 minutes, and appeared maximal after 25 mins of activating light **(Fig. 5M-Q)**. A red light-activated depolarizing channelrhodopsion ReaChR (Lin et al., 2013; Inagaki et al., 2014) also elicited a similar effect **(Figure 5, Figure Supplement 1A-C)**. Activation of ChR2 did not alter membrane abundance of the integrin component Mys (Bunch et al., 1992), or the integrin-associated protein Talin (Brown et al., 2002) **(Figure 5, Figure Supplement 1D-G’)**. In contrast, expression and activation of ChloC, a blue-light activated anion channel which hyperpolarizes cells (Wietek et al., 2017, 2014), resulted in a reduction of Smo on the plasma membrane as well as in internal compartments **(Fig. 5J-L)**. This suggests that short-term hyperpolarization generally reduces Smo levels, perhaps due to the fact that inactivated Smo is targeted for proteasome-mediated degradation following internalization, as has been reported previously (Li et al., 2012).

### Patterned expression of Rpk is necessary for maintenance of a normal compartment boundary

Our data indicate that depolarization of the membrane promotes Hh signaling, and also that expression of an activated form of Ci causes membrane depolarization, likely by increasing the expression of specific ion channels. Thus, Hh signaling and membrane depolarization appear to mutually reinforce each other just anterior to the compartment boundary. It is known that interrupting Hh signaling in this region disrupts the properties of the compartment boundary; clones of *smo* mutant cells appear to cross the boundary and occupy a position between A and P cells (Blair & Ralston, 1997; Dahmann & Basler, 2000). Therefore, we wondered whether altering *V_mem_* would impair compartment boundary function. We set out to knock down *rpk* in cells anterior to the compartment boundary using *dpp-Gal4*. Imaginal discs expressing both *dpp-Gal4* and *UAS-rpk^RNAi^* had more irregular compartment boundaries, indicating that patterned expression of Rpk in this region is necessary for maintaining a compartment boundary of normal appearance **(Fig. 6A-B’’)**.

**Figure 6.**
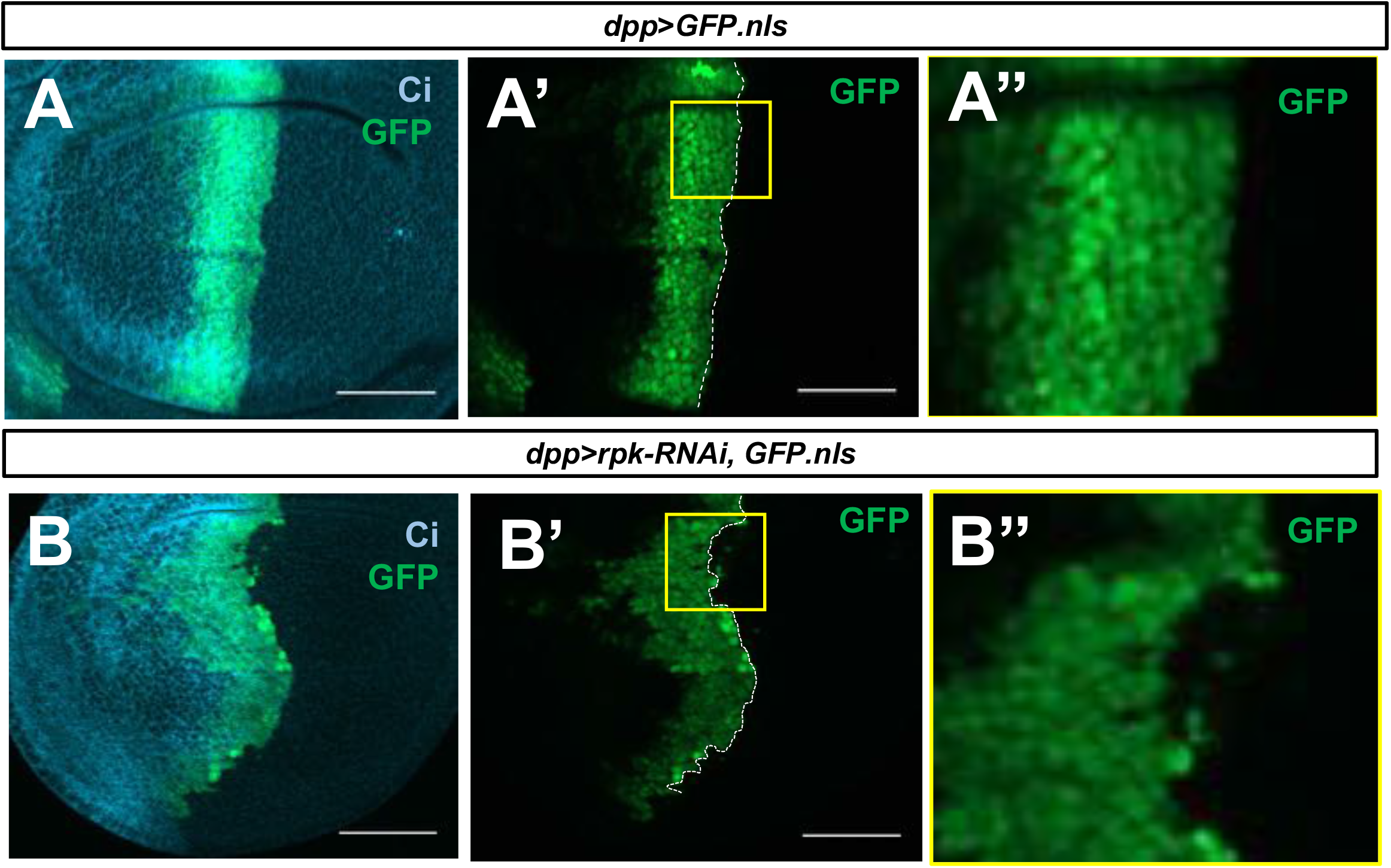
Rpk is required for compartment boundary integrity. (**A-B”**) Late L3 imaginal discs expressing *dpp-Gal4, UAS-GFP.nls* (**A-A’’’**) or *dpp-Gal4, UAS-GFP.nls, UAS-rpk-RNAi* (**B-B’’**), stained with an anti-Ci antibody (blue). Expression of rpk RNAi in the *dpp*-expressing cells anterior to the compartment boundary (**A-A’’**) results in an irregular compartment boundary compared to control discs (**B-B’’**). Boxed regions are shown at higher magnification. Scale bars are 50μM.

**Figure 5, Figure Supplement 1.**
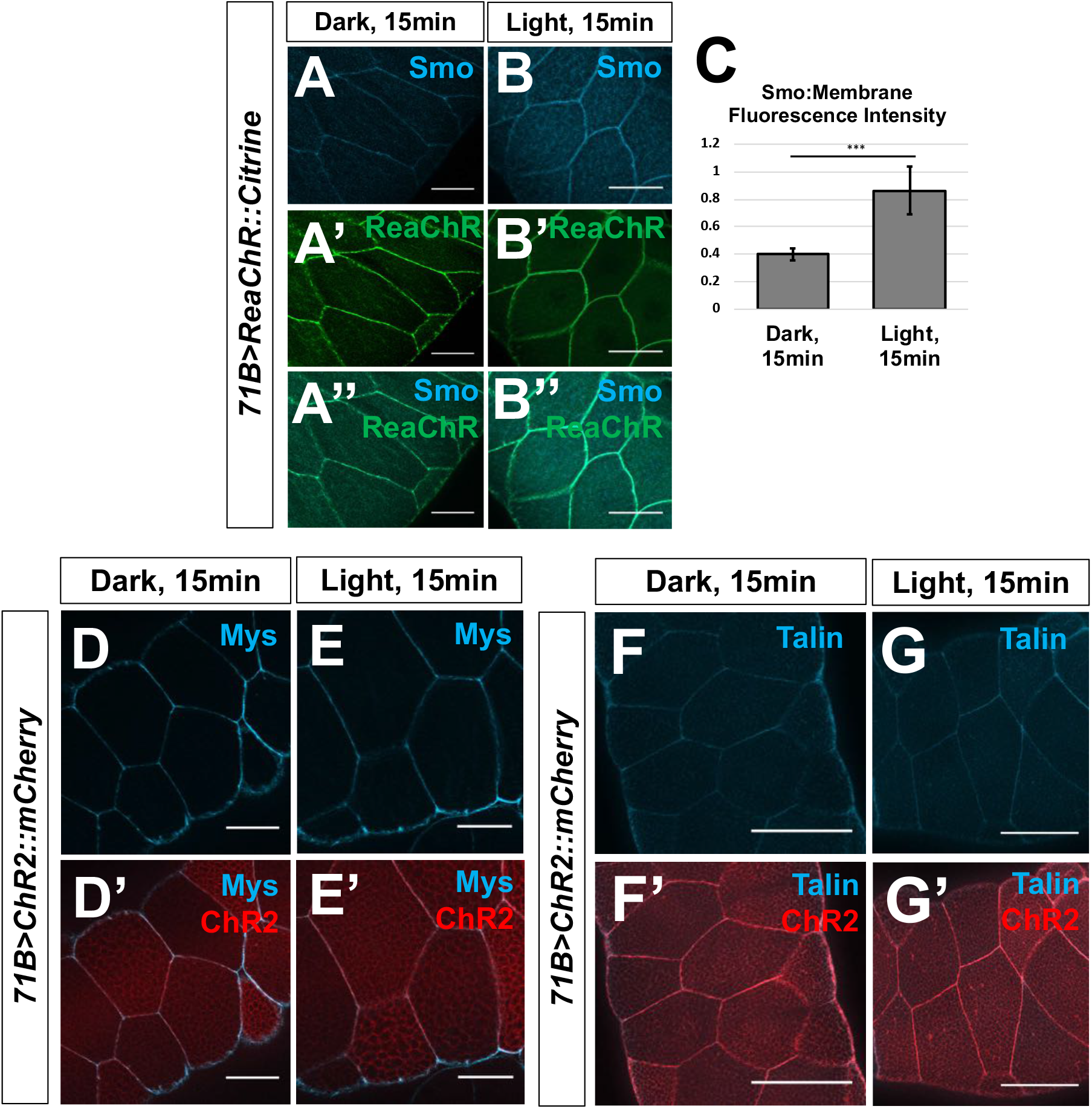
Optogenetic manipulation of *V_mem_* in Drosophila salivary glands. Salivary glands expressing the channelrhodopsin ReaChR were subjected to 15 minutes in activating light, then fixed and stained for Smo protein (**A-C**). N=9 glands, data were compared using an unpaired t test (***p<0.001). Membrane abundance of the beta integrin subunit Mys, and the integrin-associated protein Talin are not altered upon ChR2 activation (**D-G’**). Scale bars are 50μm in all panels.

### Manipulation of *V_mem_* has different effects on cell survival in the two compartments

The anteroposterior compartment boundary appears to represent not just the boundary between two lineage-restricted domains of cells, but also the boundary between two populations maintaining disparate *V_mem_*. Although A cells just anterior to the boundary appear more obviously depolarized than neighboring P cells, the pattern of DiBAC fluorescence in younger discs suggests that the zone of depolarization could be broader at earlier stages of development **(Fig. 1K-K’)**. We therefore investigated the properties of clones of cells in each of the two compartments that are more depolarized or hyperpolarized than their neighbors. We first generated discs containing clones of cells that overexpressed the bacterial sodium channel NaChBac (Ren et al., 2001; Luan et al., 2006; Nitabach et al., 2006), which would be predicted to be more depolarized than their neighbors. When clones were induced late in development, and given only 48 hours to grow in the tissue before dissection, they were recovered with equal frequency in both compartments **(Fig. 7A, B)**. However, when clones were induced at progressively earlier times in development, we observed fewer and fewer surviving clones in the P compartment **(Fig. 7A’-B**).

**Figure 7.**
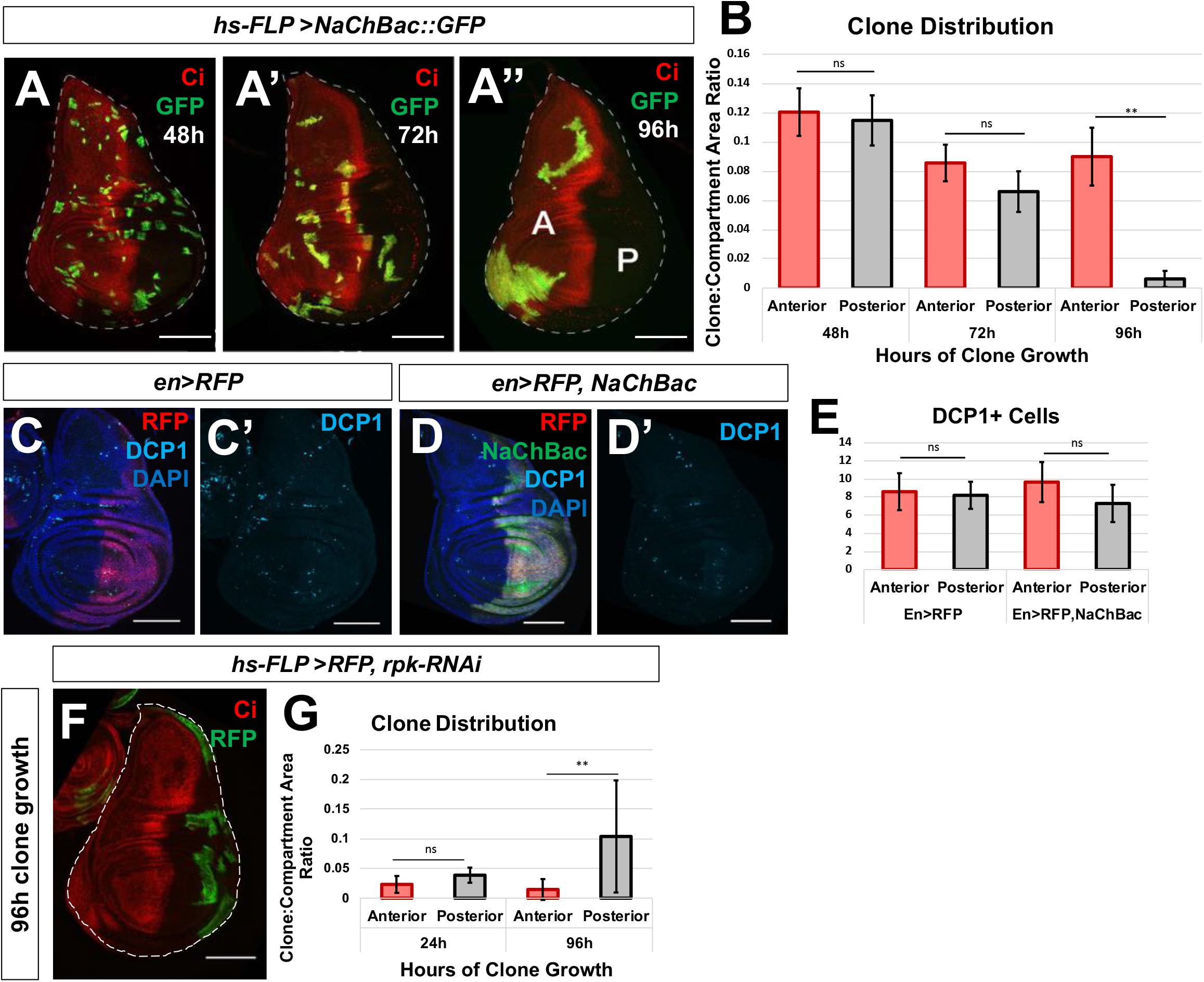
Altered membrane potential induces compartment-specific cell elimination. (**A-A’’**) Clones of cells expressing the depolarizing bacterial sodium channel NaChBac were generated 48 hours, 72 hours, and 96 hours prior to dissection. (**B**) The amount of NaChBac-expressing tissue in a compartment was quantified by measuring the ratio of the total area of GFP+ tissue to the area of the compartment, to control for the difference in size between the anterior and posterior compartments. A striking preference for clone recovery in the anterior compartment was observed in clones induced 96 hours prior to dissection, while clones induced later in development survived with relatively equal frequency between compartments, n=12 discs for each time point. Data were compared using an upaired t test (**p<0.01). (**C-E**) Elimination of clones from the posterior compartment is dependent on heterotypic interactions with wildtype cells. No difference in apoptosis was observed when NaChBac was expressed in the entire posterior compartment (**E**), n=6 discs. Data were compared using an unpaired t test. (**F-G**) Clones of cells expressing RNAi against the ENaC *rpk* are recovered preferentially in the posterior compartment, using the quantification scheme described above, n=15 discs, data were compared using an unpaired t test (**p<0.01). Clone area ratios are lower in this experiment because the heat shock exposure was decreased in order to mitigate lethality associated with widespread knockdown of *rpk*. Scale bars are 100μM in all panels.

The elimination of clones of cells over time in a context-dependent manner is reminiscent of cell competition (Morata & Ripoll, 1975; Johnston, 2009; Baker, 2017). Cell competition is a phenomenon by which cells of lower relative fitness are eliminated from a tissue due to heterotypic interactions with cells of higher fitness (Baker, 2017). We wondered whether cells expressing NaChBac were eliminated by a similar mechanism in the posterior compartment. We drove expression of NaChBac in the entire posterior compartment, forming a homotypic environment in which cells should survive if NaChBac expression is not cell-autonomously lethal. In these discs, we observed no increase in apoptosis in the NaChBac-expressing posterior compartments as compared to control discs **(Fig. 7C-E)**, suggesting that elimination of NaChBac-expressing clones in the posterior compartment is dependent on heterotypic interactions with wild-type cells.

In order to generate clones of cells that are predicted to be more hyperpolarized than their neighbors, we generated clones expressing *UAS-rpk^RNAi^*. Such clones were recovered preferentially in the P compartment, suggesting that these have lower relative fitness in the A compartment **(Fig. 7 F-G)**. Taken together, these data indicate that more depolarized cells survive better in the A compartment and that more hyperpolarized cells survive better in the P compartment.

## Discussion

In this study, we show that *V_mem_* is patterned in a spatiotemporal manner during development of the wing disc of *Drosophila*, and that it regulates Hedgehog signaling, compartment boundary maintenance, and the differential ability of cells to survive in the two compartments. First, we have shown that cells immediately anterior to the compartment boundary are relatively more depolarized than cells elsewhere in the wing pouch. This region coincides with the A cells where Hh signaling is most active, as evidenced by upregulation of Ptc. Second, we found that the expression of at least two regulators of *V_mem_*, the ENaC channel Rpk and the alpha subunit of the Na^+^/K^+^ ATPase are expressed at higher levels in this same portion of the disc. Third, by altering Hh signaling, we demonstrate that the expression of both Rpk and ATPα is increased in cells with increased Hh signaling. Fourth, by manipulating Hh signaling in the disc and using optogenetic methods in the salivary gland, we show that membrane depolarization promotes Hh signaling as assessed by increased membrane localization of Smo, and expression of the target gene Ptc. Thus, Hh-induced signaling and membrane depolarization appear to mutually reinforce each other and thus contribute to the mechanisms that maintain the segregation of A and P cells at the compartment boundary. Finally, we show that making *V_mem_* more positive is deleterious in the P compartment while decreasing it is deleterious in the A compartment.

### Developmentally regulated patterning of *V_mem_* in the wing disc

We have observed two regions of increased DiBAC fluorescence in the wing imaginal disc. Our studies have focused on the region immediately anterior to the A-P compartment boundary in the wing disc. Additionally, in the late L3 wing disc, we observe a region of increased DiBAC fluorescence in the A compartment in the vicinity of the D-V boundary. This corresponds to a “zone of non-proliferating cells” (ZNC) (O’Brochta & Bryant, 1985). Interestingly, the ZNC is different in the two compartments. In the A compartment, two rows of cells are arrested in G2 while in the P compartment, a single row of cells is arrested in G1 (Johnston & Edgar, 1998). Our observation of increased DiBAC fluorescence in the DV boundary of only the A compartment is consistent with previous reports that cells become increasingly depolarized as they traverse S-phase and enter G2 (Cone, 1969), reviewed in (Blackiston et al., 2009). In contrast, cells in G1 are thought to be more hyperpolarized. Additionally, we observed increased expression of the ENaC channel Rpk in two rows of cells at the D-V boundary in the anterior compartment (**Fig. 2A’’**), indicating that increased expression of Rpk could contribute to the depolarization observed in those cells. We note, however, that the increased DiBAC fluorescence in these cells was not entirely eliminated by exposing discs to amiloride, indicating that other factors are also likely to contribute.

While we observed a clear increase in DiBAC fluorescence anterior to the compartment boundary in the wing disc, we did not observe this in the leg disc **(Figure 1 Supplement 1B-B’)**. This could either be because the change in *V_mem_* at the compartment boundary is below that threshold of detection with DiBAC, or that the phenomenon is specific to the wing disc. In the eye-antennal disc, the region immediately anterior to the region of Hh expression is the morphogenetic furrow. Because of the change in disc architecture in this region, it is difficult to evaluate alterations in DiBAC fluorescence in unfixed preparations. We note, however, that a portion of the eye disc which represents the primordium of the ocelli has increased DiBAC fluorescence and increased Ptc expression **(Figure 1 Supplement 1A-A’)**. As more sensitive reagents for detecting changes in *V_mem_* become available, some of these issues may be clarified with greater confidence.

### How does membrane depolarization relate to Hh signaling?

Our data are consistent with a model where membrane depolarization and Hh-induced signaling mutually reinforce each other in the cells immediately anterior to the compartment boundary. Both membrane depolarization and the presence of Hh seem necessary for normal levels of activation of the Hh signaling pathway in this region; neither alone is sufficient (**Figure 8**). First, we have shown that Hh signaling promotes membrane depolarization. We have also shown that the expression of Rpk just anterior to the A-P compartment boundary is dependent upon Hh signaling. Elevated Rpk expression is not observed when a *hh^ts^* allele is shifted to the restrictive temperature, and cells become more depolarized when Hh signaling is constitutively activated through expression of the Ci3m allele. Previously published microarray data suggests that Rpk, as well as another ENaC-family channel Ppk29 are both enriched in cells that also express Ptc (Willsey et al., 2016). However, there is no antibody to assess Ppk29 expression currently. The sensitivity of the depolarization to amiloride indicates that these and other ENaC channels make an important contribution to the membrane depolarization.

**Figure 8.**
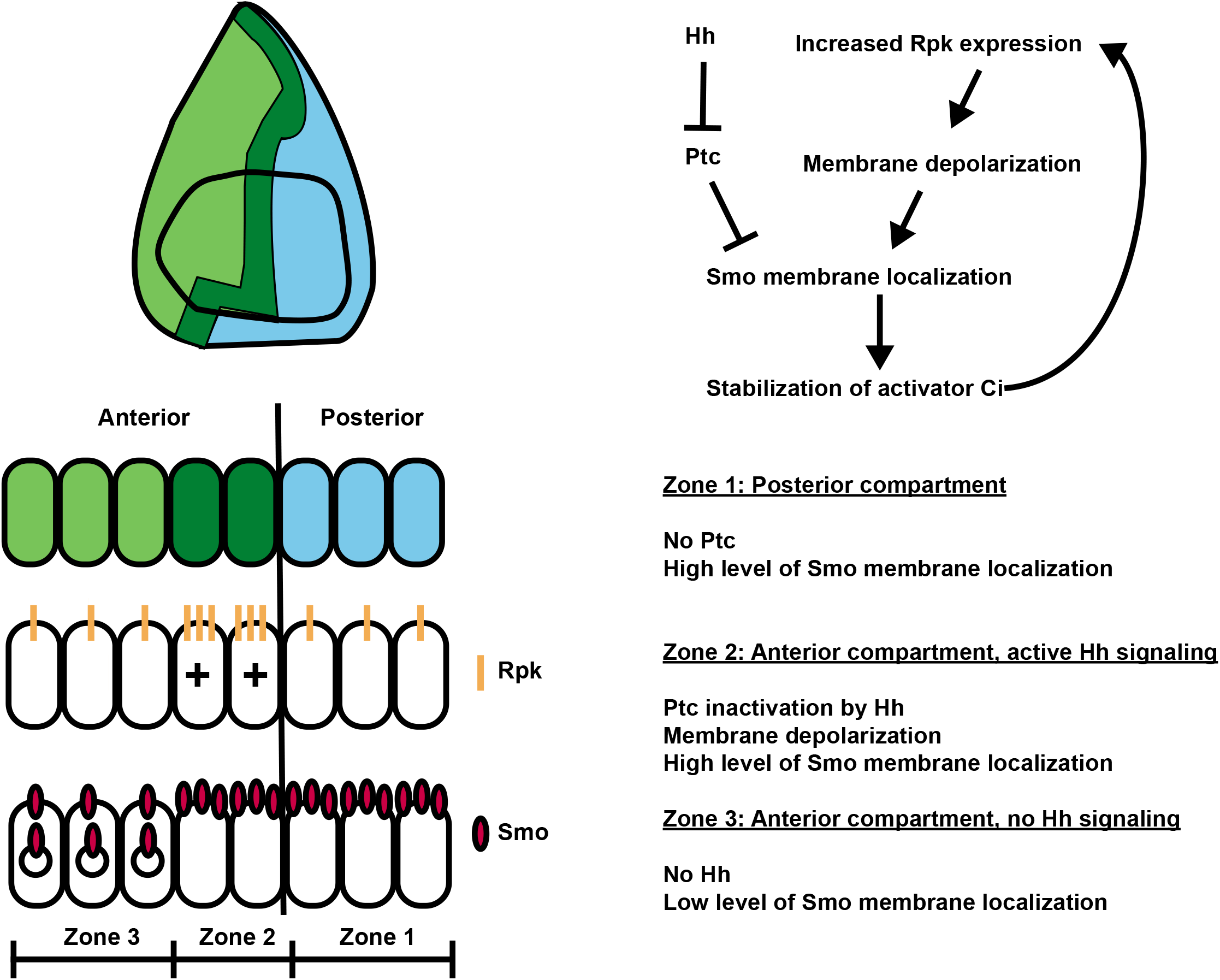
Membrane potential modulates Hh signaling at the anteroposterior compartment boundary. Cells in the posterior compartment (blue) produce the short-range morphogen Hedgehog, which is received by cells immediately anterior to the compartment boundary, in the anterior compartment (dark green). Hh signal releases Ptc-mediated inhibition of Smo, allowing for the stabilization of the transcriptional co-effector Ci, and increased expression of the ENaC channel Rpk, which depolarizes cells immediately anterior to the compartment boundary. Depolarization results in increased membrane abundance of the Hh activator Smo, and increased transduction of Hh signal. This, in turn, increases expression of Rpk, reinforcing high levels of Hh signal transduction in the region immediately anterior to the compartment boundary.

Second, we have shown that the depolarization increases Hh signaling. The early stages of Hh signaling are still incompletely understood (reviewed by (Petrov et al., 2017)). Hh is thought to bind to a complex of proteins that includes Ptc together with either Ihog or Doi. This alleviates an inhibitory effect on Smo, possibly by enabling its access to specific membrane sterols. Interestingly, it has recently been proposed that Ptc might function in its inhibitory capacity by a chemiosmotic mechanism where it functions as a Na^+^ channel (Myers et al., 2017). An early outcome of Smo activation is its localization to the membrane where its C-terminal tail becomes phosphorylated and its ubiquitlylation and internalization are prevented (Zhang et al., 2004; Li et al., 2012). By manipulating channel expression in the wing disc, and by optogenetic experiments in the salivary gland, we have shown that membrane depolarization can promote Hh signaling as assessed by increased Smo membrane localization and increased expression of the target gene Ptc. The time course of Smo activation is relatively rapid (over minutes) and is therefore unlikely to require new transcription and translation. In the P compartment, membrane Smo levels are elevated because of the complete absence of Ptc and some downstream components of the Hh signaling pathway are known to be activated (Ramirez-Weber, 2000). However, since Ci is not expressed in P cells, target gene expression is not induced. In the cells just anterior to the boundary, the partial inhibition of Ptc by Hh together with membrane depolarization seem to combine to achieve similar levels of Smo membrane localization. More anteriorly, the absence of this mutually reinforcing mechanism appears to result in Smo internalization (**Figure 8**).

Our experiments do not point to a single mechanism by which depolarization promotes Hh signaling. It is possible that depolarization results in increased Ca^2+^ levels by opening Ca^2+^ channels at the plasma membrane or by promoting release from intracellular sources (e.g. the ER or mitochondria). Indeed, there is evidence that Ca^2+^ entry into the primary cilium promotes Hh signaling, and recent work shows that targets of Sonic Hedgehog (Shh) signaling during mammalian development is augmented by Ca^2+^ influx (Delling et al., 2013; Klatt Shaw et al., 2018). A second possibility is that membrane depolarization could, by a variety of mechanisms, activate the kinases that phosphorylate the C-terminal tail of Smo and maintain it at the plasma membrane in an activated state. Depolarization could also impact electrostatic interactions at the membrane that make the localization of Smo at the membrane more favorable. Finally, since a chemiosmotic mechanism has been hypothesized to drive Smo inhibition by Ptc (Myers et al. 2017), depolarization could oppose an incoming Na^+^ current, which would be predicted to reduce the inhibitory effect of Ptc on Smo. Importantly, all these mechanisms are not mutually exclusive and their roles in Hh signaling are avenues for future research.

### A compartment-specific cell elimination phenomenon could reinforce compartmentalization of the disc

Our data indicate that the regulation of *V_mem_* could contribute to compartmentalization of the wing disc in at least two different ways. First, Hh signaling, which is strongest in the cells just anterior to the compartment boundary, is augmented by membrane depolarization. Previous work has shown that A cells at this location that are deficient in transducing a Hh signal fail to remain in the A compartment and can cross the compartment boundary (Blair & Ralston, 1997; Dahmann & Basler, 2000). Consistent with this we have found that expression of *rpk-RNAi* in these cells results in an irregular compartment boundary, indicating that there is increased intermingling of A and P cells.

We have also found a second mechanism that could contribute to the segregation of A and P cells. Clones of cells that are more depolarized than their neighbors tend to survive preferentially in the A compartment while clones of hyperpolarized cells tend to survive preferentially in the P compartment. Thus, single cells from each compartment that enter the other compartment could be eliminated.

The compartment-specific mechanism of cell elimination that we have uncovered has some similarities to the group of phenomena referred to as cell competition that was first described for clones of cells that were heterozygous for a *Minute* mutation (Morata & Ripoll, 1975). A gene that has been implicated in cell competition, *flower* (*fwe*) encodes a Ca^2+^ channel (Yao et al., 2009; Rhiner et al., 2010), but compartment specificity in the behavior of *fwe* clones has not been described thus far, and cells expressing a specific isoform of *fwe* are eliminated in both compartments (Rhiner et al., 2010). Thus, there is currently no simple way to connect the compartment-specific clone elimination phenomenon we have observed to existing models of cell competition. The relationship between clone elimination caused by changes in *V_mem_* and other types of cell competition therefor represent a fertile area for future research.

It is now generally accepted that both cell-cell signaling and mechanical forces have important roles in cell fate specification and morphogenesis. Our work here adds to a growing body of literature suggesting that changes in *V_mem_*, a relatively understudied parameter, may also have important roles in development. Integrating such biophysical inputs with information about gene expression and gene regulation will lead to a more holistic understanding of development and morphogenesis.

## Acknowledgements

We thank the members of the Hariharan lab, Ehud Isacoff, Diana Bautista, David Bilder, Kristin Scott, Henk Roelink, Michael Levin, Evan Miller, and Julia Lazzari-Dean for helpful discussions and feedback. We also thank Diana Bautista, David Bilder, Dan Kiehart, and Kristin Scott for reagents and fly stocks. This work was funded by NIH grant R35 GM122490 and IKH is a Research Professor of the American Cancer Society (RP-16238-06-COUN).

## Materials and Methods

### Drosophila strains and husbandry

Animals were raised on standard medium as used by the Bloomington *Drosophila* Stock Center. All animals were raised at 25ºC, except *ap>Rpk-RNAi, dpp>Rpk-RNAi*, and *ap>Irk1-RNAi* crosses, which were raised at 18ºC to reduce lethality.

Stocks used in this study include: *hsFLP;; act<stop<Gal4, UAS-RFP/S-T, yw;ap-Gal4/Cy0; TM2/TM6B*, *;;ap-Gal4,; ptc>RFP;,;; dpp>GFP, w;;UAS-Ci3M,;; 71B-Gal4*, and *w;UAS-Kir2.1; UAS-Kir2.1/TM6B* (from Kristin Scott, UC Berkeley, USA).

Stocks obtained from the Bloomington Stock Center (Bloomington, IN, USA) include *UAS-ArcLight* (BL:51056)*, UAS-Rpk-RNAi*; (BL:39053, 25847)*, UAS-Irk1-RNAi* (BL:25823)*, UAS-NaChBac*; (BL:9466)*, UAS-ChR2::mCherry*; (BL:28995)*, UAS-ReaChR::Citrine*; (BL:53741)*, UAS-ChloC::tdTom* (BL:76328)*, en-Gal4*; (BL:6356).

### Live imaging and optogenetics

Larvae were washed with 70% EtOH and PBS prior to dissection. Live tissue was dissected in Schneider’s media (#21720001, Gibco), and care was taken to not damage or stretch tissue. For DiBAC staining, imaginal discs were incubated in 1.9μM DiBAC_4_(3) (bis-(1,3-dibutylbarbituric acid) trimethine oxonol; DiBAC_4_(3); Molecular Probes)) in Schneider’s media for 10 minutes with gentle rotation. A small amount media was used to mount the discs, such that addition of a coverslip did not destroy the tissue, and discs were imaged right away. 100μM amiloride (#A7419, Sigma-Aldrich) and 100μM ouabain (#O3125, Sigma-Aldrich) were added to DiBAC solution for pharmacology experiments. Discs imaged in FM4-64 dye (#T13320, ThermoFisher) were incubated in 9μM FM4-64, and imaged without washing, to preserve staining of the cell membrane. Discs expressing ArcLight were dissected, mounted, and imaged in Schneider’s.

For optogenetics experiments, carcasses were dissected and cleaned (fat body removed) in Schneider’s medium. Carcasses were loaded onto a glass cover slip in a large drop of Schneider’s, and either kept in the dark (control condition) or exposed to activating light (480nm for ChR2 and ChloC experiments, 647nm for ReaChR experiments). After exposure, carcasses were immediately fixed and prepared for immunohistochemistry.

### Immunohistochemistry

Imaginal discs were dissected in phosphate buffered saline, fixed for 20min in 4% PFA at room temperature, permeabilized in phosphate buffered saline with 0.1% Triton X-100, and blocked in 10% Normal Goat Serum. Primary antibodies used were: rat anti-Ci (1:10 Developmental Studies Hybridoma Bank, DSHB), mouse anti-Smo (1:10, DSHB), guinea-pig anti-Rpk (1:500) (gift from Dan Kiehart) mouse anti-ATPalpha (1:100, DSHB), mouse anti-Ptc (1:50, DSHB), rabbit anti-cleaved-DCP1 (Asp216, 1:250; Cell Signaling Technology), and rabbit anti-GFP (#TP401, 1:500; Torrey Pines Biolabs). Secondary antibodies used were: goat anti-rabbit 488 (#A32731; Invitrogen), goat anti-mouse 488 (#A32723; Invitrogen), goat anti-mouse 555 (#A32727; Invitrogen), goat anti-rat 555 (#A-21434; Invitrogen), goat anti-mouse 647 (#A32728; Invitrogen), goat anti-rabbit 647 (#A32733; Invitrogen), and goat anti-rat 647 (#A-21247; Invitrogen). Nuclei were stained with DAPI (1:1000, Cell Signaling). Samples were imaged on a Zeiss Axio Imager.M2 with Apotome.2.

### Quantification and statistical analysis

Fluorescence intensity and area measurements were recorded using FIJI software (NIH, Bethesda, USA). *P* values were obtained using ANOVA and unpaired Student’s *t* tests (Graphpad). Error bars in all graphs are standard deviation. *P* value significance <0.001: ***; 0.001 to 0.01: **; 0.01 to 0.05: *; >0.05: not significant.

### Mosaic Tissue Generation

Experiments using heat shock-controlled FLPase and *act≪Gal4* were maintained at 25C. To generate clones overexpressing NaChBac, *ywhs-FLP;;Act≪Gal4,UAS-RFP/S-T* virgin females were crossed to *yw;;UAS-NaChBac* (BL:9466) males. Crosses were set up in cages, and females were allowed to lay eggs on grape plates with yeast paste for 8 hour increments. L1 larvae were picked and seeded into vials at a density of 50 larvae/vial. Vials were subjected to a 10 minute heat shock in a 37ºC water bath 96, 72, and 48 hours prior to dissection. To generate clones expressing RNAi against Rpk, *hs-FLP;;act<STOP<UAS-RFP/S-T* virgin females were crossed to *yw;;UAS-Rpk-RNAi* (BL:39053, BL:25847) males. Larvae were collected as described above, but vials were subjected to five minute heat shocks in order to mitigate lethality associated with broad expression of Rpk-RNAi. To generate clones expressing Ci3m, *hs-FLP;;act<STOP<UAS-RFP/S-T* virgin females were crossed to *UAS-Ci3m* males. Larvae were collected as described above, and vials were subjected to a 10 minute heat shock in a 37ºC water bath 48 hours before dissection and live imaging in DiBAC.

